# Accelerating Neuron Reconstruction with PATHFINDER

**DOI:** 10.1101/2025.05.16.654254

**Authors:** Michał Januszewski, Thomas Templier, Kenneth Hayworth, David Peale, Harald Hess

## Abstract

Comprehensive mapping of neural connections is essential for understanding brain function. Existing automated methods for connectome reconstruction from high-resolution images of brain tissue introduce errors that require extensive and time-consuming manual correction, a critical bottleneck in the field. To address this, we developed PATHFINDER, an AI system that segments volumetric image data, identifies potential ways to assemble neuron fragments, and evaluates the plausibility of resulting shapes to reconstruct complete neurons. Using a dataset of all axons in an IBEAM-mSEM volume of mouse cortex, we show that PATHFINDER reduces the error rate in axon reconstruction by an order of magnitude over previous state of the art, leading to an improvement in proofreading throughput of up to 84× relative to prior estimates in the context of a whole mouse brain. By drastically reducing the manual e”ort required for analysis, this advance unlocks the potential for both large-scale connectome mapping and routine investigation of smaller volumes.

## 1. Introduction

Connectomics, the study of neural connections, requires comprehensive tracing of individual neurons within high-resolution, three-dimensional images of brain tissue acquired at nanometer resolution. The estimated rate of human tracing (∼ 1 mm /h with optimized tools (Boergens et al., 2017)) renders manual reconstruction impractical, necessitating the use of automated methods. Substantial progress in automated tracing (Januszewski et al., 2018; Macrina et al., 2021; Sheridan et al., 2023) has been achieved in recent years, with analyzed volumes reaching the cubic millimeter scale (MICrONS Consortium; Shapson-Coe et al., 2024). While algorithms are now capable of generating reconstructions with error-free path lengths on the order of a millimeter, this is still insufficient for analyzing complete neuronal processes, which often extend over distances at least an order of magnitude greater. Consequently, investigations of the graph structure of connectomes require manual correction of errors introduced by the automated methods. Recent large-scale reconstruction projects in Drosophila (Dorkenwald et al., 2024; Scheffer et al., 2020; Takemura et al., 2024) required tens of person-years of manual proofreading effort for a single insect brain. This constitutes a significant bottleneck for connectomic endeavors, particularly as the field aims to scale up to mapping the whole brains of small mammals (Abbott et al., 2020), four orders of magnitude larger than that of the fruit fly. Jefferis et al. (2023) estimated that if current reconstruction techniques were to be used, proofreading a mouse brain would cost billions of dollars, consuming over 90% of the total budget of the project. Further progress in automated tracing is therefore critically needed.

Automated neuron reconstruction in connectomics is typically a two-step process that first segments volumetric images into a „base segmentation” of supervoxels, and then agglomerates these into larger neuron fragments. There are three categories of connectome-relevant errors that a neuron reconstruction system can make: *splits* (more than one segment assigned to the same neuron), *mergers* (a segment covering more than one neuron), and *omissions* (a part of the image representing a neuron not covered by a segment). Supervoxels are treated as atomic units in downstream processing, but merge errors within them can be reduced through ensembling techniques such as oversegmentation consensus (OC) (Januszewski et al., 2018), at the expense of increased splits. As long as the false merge rate of the supervoxels is sufficiently low, the accuracy of the final reconstruction in this two-step approach hinges on the performance of the agglomeration system.

Existing state-of-the-art methods leverage the base segmentation model for agglomeration, either by directly reusing its predictions (mean-affinity agglomeration (Lee et al., 2017)) or by recomputing them for pairs of neighboring supervoxels (Januszewski et al., 2018). This approach is limited in a number of ways. First, base segmentation errors (e.g., omissions due to image irregularities) currently restrict the analysis to an incomplete set of true positive supervoxel pairs. Pairs that should be considered are missed if they are separated by unsegmented regions or incorrectly segmented fragments. Second, the field of view of the base segmentation model is optimized for voxel-level 3d image processing, limited by GPU memory, and too small to take advantage of all the morphological cues provided by the complete supervoxels which can extend over many microns. Finally, with agglomeration scores computed independently in a pairwise manner, locally optimal decisions can sometimes lead to a suboptimal global result.

To overcome these limitations, we introduce PATHFINDER (Precise Analysis, Tracing and High Fidelity Interpretation of Neuronal Data for Exhaustive Reconstruction), a novel multi-stage AI system using specialized and complementary models to process data across progressively larger spatial scales (Figure 1D). The models are:

**Figure 1.**
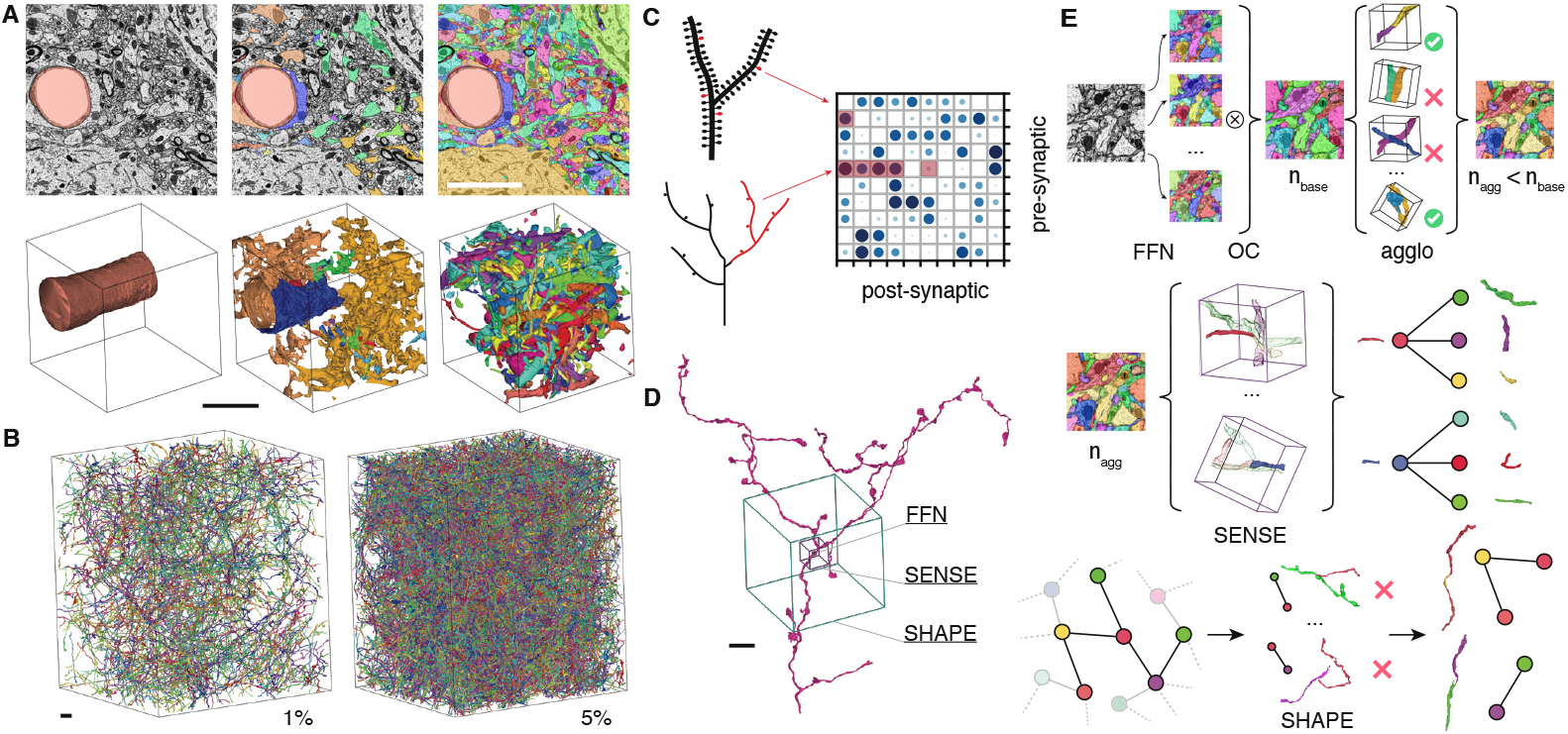
**A**: Sequential segmentation. Columns left to right: blood vessels, glia, and neurons. Top row: 2D cross sections. Bottom row: 3D renderings (somas omitted for clarity). **B**: Density of axons within the volume. Two random subsets of all unmyelinated axons in the volume are shown, representing 1% (left) and 5% (right) of the total unmyelinated axon path length (4.3 m) within the volume. **C**: Impact of typical dendrite (top) and axon (bottom) splits on connectome matrix (entries missing from the true connectivity matrix because of the split errors are highlighted). **D**: Comparison of the field of view (FOV) of the stages of the PATHFINDER system: FFN (0.5 µm, black), SENSE (4 µm, purple), and SHAPE (20 µm, teal). **E**: System-level overview of PATHFINDER. Top row: FFNs generate multiple segmentations of the EM volume, which are then combined into a base segmentation through oversegmentation consensus (OC), reducing merge errors at the cost of a moderate increase in splits. Pairs of segments are evaluated by the FFN again and agglomerated when possible, reducing split errors and the total number of segments. Middle row: SENSE models identify more agglomeration candidates for every segment, forming an overcomplete agglomeration graph. Bottom row: motifs from this agglomeration graph are evaluated by the SHAPE model and edges corresponding to invalid segment combinations are removed. Remaining connected components correspond to larger neurites assembled from the segments. All scale bars 5 µm.

- Flood-Filling Networks (FFN) v1.5: an optimized small field-of-view segmentation model and updated training and inference protocols (Januszewski et al., 2018) to produce large supervoxels with low merge error rates,
- SinglE Neuron Segment Expander (SENSE): a wide field-of-view model to identify a comprehensive set of agglomeration candidates,
- Scoring Hypothesis Assemblies using Plausibility Evaluation (SHAPE): a model focused on morphological features to evaluate the plausibility of the shape of a neuron fragment represented by one or more supervoxels.

Figure 1E illustrates the PATHFINDER workflow. First, FFNs build the base segmentation (Figure 1A). Next, an ensemble of multi-resolution SENSE models defines the space of possible agglomerations. Finally, the SHAPE model guides an efficient combinatorial search through this space to find optimal agglomerations, resulting in reconstructed neurons. We demonstrate the effectiveness of this approach by tracing all axons in a 80 × 85 × 99 µm^3^ volume of mouse cortex tissue (Figure 1B), resulting in a → 84 × reduction in estimated proofreading cost relative to the optimistic variant of the projections of Jefferis et al. (2023). This makes large-scale connectome mapping projects such as the whole mouse brain significantly more economically feasible and paves the way towards routine analysis of smaller volumes.

## 2. Results

### Image volume and ground truth

We used an IBEAM (Ion Beam Etching and Milling) (Hayworth et al., 2020; Templier, 2018) setup connected to a multibeam scanning electron microscope (Eberle and Zeidler, 2018) to image a block of mouse cortex tissue at a voxel size of 8×8×8 nm^3^, from which we selected a 80.4×84.8×99.2 µm^3^ cuboid (∼1.32 TB of data) for further analysis.

We obtained a reconstruction of the entire volume and used a neural network-based tissue classification model (Shapson-Coe et al., 2024) to assign processes to different types: axons, dendrites, or others (Methods). Following this identification, all 4.45 m of axons contained within the volume, as well as all dendrite trunks, were manually proofread to form a reference reconstruction (Supplementary). This morphologically complete reference was used to compute whole-volume topological metrics for evaluating segmentation quality (Table 1).

**Table 1.** Axon quality evaluation by reconstruction stage.

| STAGE | NERL | SPLITS | MERGERS |
| --- | --- | --- | --- |
| FFN v1.0 (single) | 68.5% | 44,417 | 1,282 |
| FFN v1.0 (base) | 56.2% | 79,230 | 200 |
| FFN v1.0 (agglo) | 76.6% | 33,835 | 347 |
| FFN v1.5 (single) | 78.5% | 28,207 | 196 |
| FFN v1.5 (base) | 66.1% | 50,981 | 20 |
| FFN v1.5 (agglo) | 82.0% | 23,085 | 33 |
| PATHFINDER | 94.2% | 2,862 | 702 |

Smaller neurite calibers increase the difficulty of reconstruction by humans and automated methods. Axons generally have a smaller diameter and a greater length than dendrites. Also, the impact of tracing errors on connectome accuracy is much greater for axons compared to dendrites (Figure 1C). To demonstrate the effectiveness of PATHFINDER we therefore focused our analysis on the more challenging axons only.

### Base segmentation

We modified the previously proposed FFN workflow (Januszewski et al., 2018) to create a base segmentation of the volume. Briefly, we enhanced the neural network capacity by incorporating additional residual modules to increase its depth while keeping the network architecture largely fixed (Methods). We also modified the training procedure by simplifying seed point sampling to match the inference-time process and used imagery downsampled 2 × in all dimensions to 16×16×16 nm^3^ per voxel as the input, reducing processing cost by a factor of eight.

We found the initial FFN base segmentation to exhibit too many merge errors (Table 1). Merge errors can be reduced with a previously introduced oversegmentation consensus (OC) technique (Januszewski et al., 2018), which compares several candidate segmentations and outputs a new one in which two voxels are assigned to the same segment if and only if they belong to the same segment in every candidate segmentation. We applied OC with inverse seed ordering and with additional FFN checkpoints (snapshots of network weights saved at different points of the training phase) to reduce the merge rate by 9.9 ×, at a cost of a 1.8× increase of split rates. The segmentation was then agglomerated with FFN agglomeration, reducing split rates by a factor of 2.2× (Methods). FFN agglomeration evaluates pairs of nearby segments by performing two separate FFN segmentations within a small EM subvolume, each seeded from one of the segments. The similarity between these two outputs yields an agglomeration score for the segment pair.

### Agglomeration candidate generation

We observed that the effectiveness of current agglomeration methods is limited by the incomplete set of segment pairs considered for merging. FFN agglomeration does not require segments to be directly adjacent in voxel space and utilizes spatial proximity for pair selection, but still misses true agglomeration pairs. Relaxing geometric constraints would improve recall, but the computational cost of pair scoring is high due to quadratic scaling.

To address this limitation, we propose a novel approach that does not require a separate pair identification step prior to running the scoring model. We introduce SENSE, a model which determines the likely continuation of a segmented fragment into surrounding unlabeled electron microscopy voxels by analyzing the fragment and its local image context (Figure 2B). While conceptually similar to the FFN, SENSE employs a U-net architecture, utilizes fixed-size segment prompts (here, spherical with *r*=16 vx), and processes data within a larger field of view (*d*=2-4 µm) compared to the FFN base segmentation model (*d*=0.5 µm, Figure 1D). Unlike previous work (Schmidt et al., 2024), SENSE processes raw images directly without relying on any geometric transformations or directional biases.

**Figure 2.**
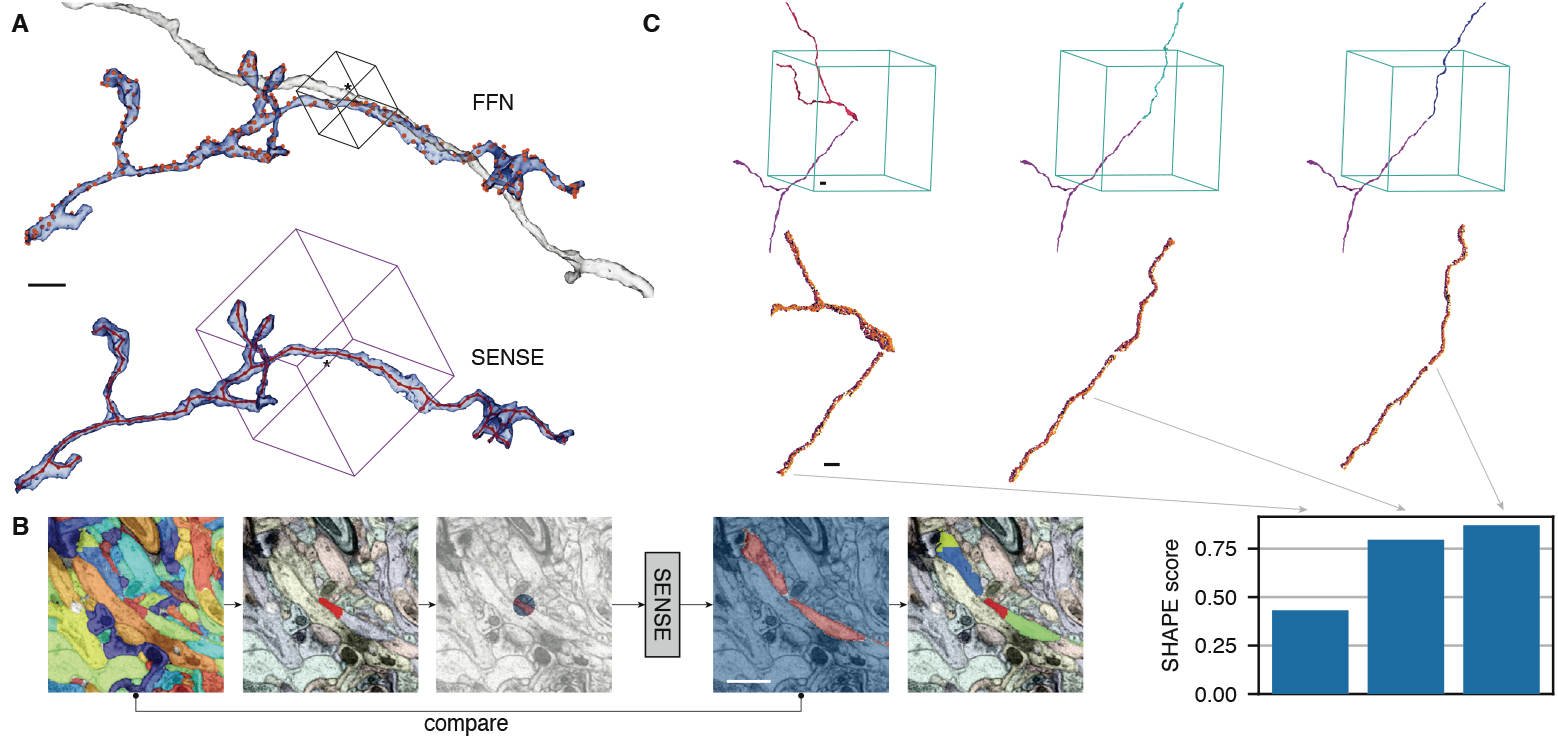
**A**: Agglomeration candidate selection for an axon fragment (blue). Top: for FFN, points of maximal proximity (orange) of neighboring segments are used. A single agglomeration partner is shown (grey) for the point marked with *. Bottom: SENSE is evaluated at every skeleton node (red). The FOV for the node marked with * is shown (purple cube). **B**: A single supervoxel from an existing segmentation is selected, converted into a „prompt” and expanded to the full FOV by SENSE. The results are compared with the initial segmentation to determine agglomeration candidates (highlighted segments). **C**: Scoring SENSE agglomeration hypotheses with SHAPE models. Candidate agglomeration pair meshes are assembled (top row), truncated to a central spherical part of radius 10 µm, and points are sampled from the surfaces (middle row), which are then scored by SHAPE (bottom bar plot). The highest scoring (rightmost) variant is the correct combination. All scale bars 1 µm.

We used a set of manually proofread axons to train SENSE models with 16× 16 × 16 nm^3^ and 32 × 32 × 32 nm^3^-voxel input data (Methods). We then reduced the base segmentation to a skeleton representation and performed inference with the SENSE models, centering queries at skeleton nodes (Figure 2A). Merge candidates for each query segment were generated by comparing the model outputs with the base segmentation by voxel overlap, potentially resulting in multiple candidates from a single query (Figure 2B). In contrast to the quadratic-complexity pair agglomeration procedure used in the original FFN workflow, SENSE inference scales linearly with the number of segments and is therefore computationally more efficient.

We applied the SENSE models in a high-recall configuration intended to capture all true positive segment pairs. This however inherently increases the number of false positives (merge errors, Figure 4A). Therefore, the primary role of SENSE within PATHFINDER is to generate a comprehensive superset of candidate segment pairs for subsequent filtering, rather than to directly improve reconstruction.

### Neuron shape plausibility evaluation

Experienced human annotators intuitively understand neuronal morphology, enabling them to quickly identify potential errors in reconstructions when looking at large branches or complete neurons. Nevertheless, well-trained annotators can still make errors when tracing at a high level of detail and focused on small subvolumes of data, particularly in regions with ambiguous imagery. These errors look locally plausible but result in improbable shapes when viewed from a wider perspective.

We hypothesized that a model equipped with a larger FOV would not make such errors and could serve as an effective filter for the agglomeration candidates generated by SENSE. Representing a neuronal fragment within a large subvolume using a dense voxel grid is inefficient, because individual neurons typically occupy a small fraction of the total volume and the expected fill rate scales as 1 / *d*^2^ as a function of subvolume diameter *d*. To address this, we adopted a point set representation, sampling points from the mesh faces of the segments (Figure 2C). A similar strategy for data representation has been proposed in the context of an automated proofreading workflow aimed at correcting merge errors by clustering (Troidl et al., 2024).

We trained SHAPE, a variant of the PointNeXT model (Qian et al., 2022) with a spherical *r* = 10 µm field of view, to differentiate between correct and incorrect neuronal shapes (Figure 2D), utilizing a superset of the axons previously used for training SENSE models (Methods). Correct shapes were taken directly from the training set, while incorrect shapes were generated by merging axons with other segments identified by SENSE operating in the high-recall regime. This approach ensured the model’s relevance to practically encountered scenarios.

### Combinatorial search for agglomeration

We filtered SENSE-generated agglomeration candidate segment pairs with SHAPE, which enhanced reconstruction quality (Figure 4A) by reducing false mergers. However, the SHAPE model can evaluate arbitrary objects and is not restricted to pairs of supervoxels. We therefore used SHAPE to score progressively larger subgraphs corresponding to combinations of segments within local subvolumes (Figure 3), identifying agglomerates likely to cause merge errors. We then employed an iterative optimization and global reconciliation process to identify graph edges that need to be removed to prevent these merge errors while minimizing the introduction of new splits (Methods).

**Figure 3.**
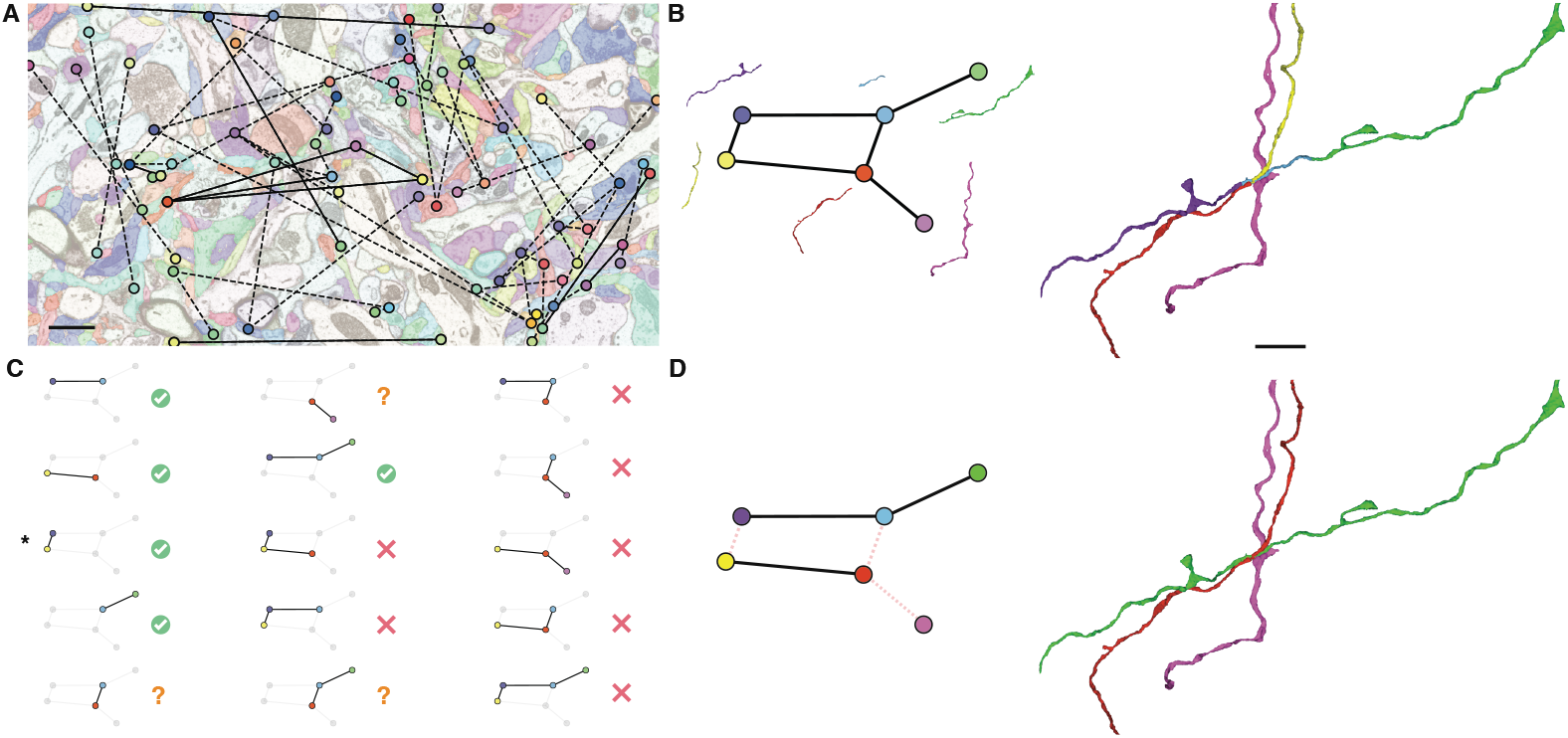
Agglomeration through combinatorial search with SHAPE models. **A**: An overcomplete (i.e., likely to lead to merge errors) proposal agglomeration graph is formed using SENSE predictions (dashed edges), and filtered by pairwise SHAPE scores (solid edges). Pairs within 1 µm of the shown section in Z are displayed. Scale bar 1 µm. **B**: A single connected component of that graph contained within a *r*=10 µm spherical FOV (left) and its corresponding segments (right). Scale bar 3 µm. **C**: Connected subgraphs of the component scored with SHAPE. Green tick indicates scores > 0.9, red cross < 0.1, question mark 0.1-0.9. One of the subgraphs marked with *⋆* is incorrectly scored as morphologically plausible. This evaluation is later corrected as other combinations involving the motif are negatively scored. **D**: Edges (dashed red) are removed from the connected component so that no subgraphs negatively evaluated by SHAPE remain (left), which results in a correct agglomeration (right).

**Figure 4.**
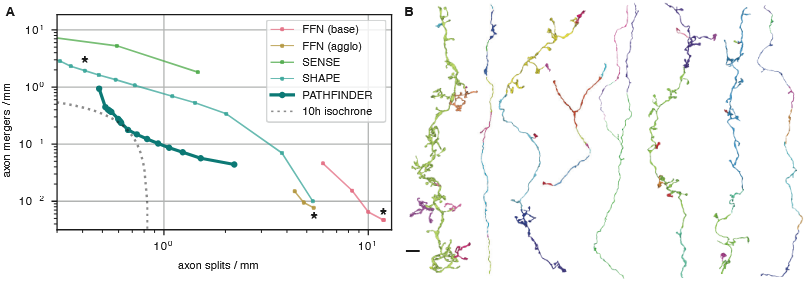
Evaluation of axon reconstruction quality over all unmyelinated axons in the volume. **A**: Split-merge trade-off for various stages of reconstruction. FFN (base) (nodes left to right): single inference pass, seed order OC, OC with two additional checkpoints. FFN (agglo): consensus using one, two, and three checkpoints. SENSE: results as a function of varying the overlap fraction threshold for partner segments. SHAPE: filtering agglomeration decisions as a function of SHAPE scores for binary decisions. PATHFINDER: Pareto optimal reconstructions with combinatorial search of graphs of up to four segments, with SHAPE score decision threshold of 0.5 and using SHAPE or SENSE scores to prioritize edge filtering. marks settings selected for use in the following stages of the PATHFINDER pipeline. Dotted line indicates split and merge rates requiring an estimated 10 h of proofreading time to correct all errors (Supplementary). **B**: Sample PATHFINDER reconstructions for different axon morphologies. Colors indicate segments after FFN agglomeration. Scale bar 5 µm.

Evaluating all subgraphs is, in the limit, a process of exponential complexity, which could quickly become unmanageable even with the limited 10 µm FOV of the SHAPE model. Our approach strategically leverages two key techniques to avoid this combinatorial explosion. First, we initiate the process with the relatively large supervoxels generated by the FFN, significantly reducing the total number of segments to consider per FOV. Second, we employ SENSE models for prefiltering, constraining the search space and eliminating unlikely combinations early. This dual approach is crucial for rendering the problem tractable, enabling efficient and accurate analysis.

The combined approach integrating SENSE, SHAPE, and the search policy to agglomerate supervoxels produced by the FFN v1.5, constitutes the PATHFINDER system. Evaluating PATHFINDER revealed improvements in segmentation quality relative to pairwise filtering with SHAPE, demonstrating the effectiveness of considering higher-order graph motifs for agglomeration (Figure 4A). The segmentation variant optimized to minimize the total number of axon split and merge errors showed a normalized expected run length (nERL) of 94.2% (Table 1). nERL, defined as ERL divided by the maximum possible ERL value for a given ground truth set, provides a robust measure of overall quality. Prior work analyzing data on a similar spatial scale achieved a global (i.e., including axons, dendrites, and cell bodies) nERL of only 52% (Januszewski et al., 2018) and required two orders of magnitude more proofreading time for volumes of comparable size (Januszewski et al., 2018; Motta et al., 2019).

## 3. Conclusions

PATHFINDER’s nERL of 94.2% for axons in an IBEAM-mSEM volume of mouse cortex translates to an estimated proofreading throughput of up to 67,200 µm^3^ / h for exhaustive axon reconstruction (Supplementary) — an 84-fold increase over previous „optimistic” estimates in the context of a potential whole mouse brain project (Jefferis et al., 2023). The low number of errors in the PATHFINDER reconstruction not only reduces the manual effort needed for error correction but should also make the purely automated results more useful before proofreading is applied. Notably, existing large-scale *proofread* connectomes (Dorkenwald et al., 2024; Scheffer et al., 2020) have demonstrated considerable practical value without eliminating all reconstruction errors and despite less than fully complete connectivity matrices.

Unlike previous approaches that aimed to emulate aspects of human proofreading (Schmidt et al., 2024) or fix errors through heuristics (Celii et al., 2025), our objective is to directly maximize the accuracy of automated reconstruction through a complete, multi-stage optimized system. The segmentations produced by PATHFINDER should therefore be complementary to and compatible with „auto-proofreading” methods. We do not mimic human decisions, relying instead on sparse yet fully traced neurites for training and the possibility to explore many potential solutions at inference time. These ground truth reconstructions are currently assembled manually, but they could potentially be obtained automatically using sparse labeling techniques and correlated light and electron microscopy (CLEM) (Drawitsch et al., 2018; Fang et al., 2018) or LICONN (Tavakoli et al., 2024), further enhancing the scalability of our approach and decoupling it from manual tracing.

Relative to recent work (Schmidt et al., 2024), the PATHFINDER reconstruction shows simultaneous improvement in split and merge error rates by an order of magnitude. By evaluating our method on *all axons* within the volume, we avoid previously raised concerns about biases in evaluation set selection (Schmidt et al., 2024). Our evaluation also encompasses a total path length two orders of magnitude larger than previous efforts (Januszewski et al., 2018; Schmidt et al., 2024), ensuring statistically robust results.

We anticipate further improvements with additional data and larger volumes, enabling the extension of the models to wider fields of view. Future work could also explore incorporating image features for enhanced accuracy, leveraging self-supervised pretraining to maximize utilization of available ground truth annotations, and employing equivariant representations for improved inference efficiency.

Historically, the ability to image and reconstruct connectomic volumes has far surpassed the capacity to proofread and analyze the resulting data. „Frontier” volumes at the 1 mm^3^ scale (MICrONS Consortium; Shapson-Coe et al., 2024) have so far remained economically infeasible to comprehensively reconstruct, necessitating targeted manual correction of specific cells. PATHFINDER marks a turning point where the once-dominant cost of proofreading becomes comparable to the cost of image acquisition and processing at the scale of a whole mouse brain (Jefferis et al., 2023). This advance paves the way for significant increases in research throughput, enabling the analysis of larger datasets and ultimately accelerating our understanding of the brain.

## Acknowledgments

We thank Sven Dorkenwald, Viren Jain, Mariela Petkova, and Franz Rieger for comments on the manuscript.

## Author contributions

MJ developed the PATHFINDER system, applied it to the EM volume, and evaluated and visualized the results. TT, KH, DP, and HH developed the IBEAM-mSEM system and acquired the mouse cortex volume. TT, KH, and MJ registered and aligned the EM volume. MJ wrote the paper, with inputs from all authors. All authors read and approved the manuscript.

## Data and materials availability

The dataset and code will be released upon publication.

## Supplementary materials

Methods

Supplementary Text

Figs. S1 to S3

Tables S1 to S2

**Figure S1.**
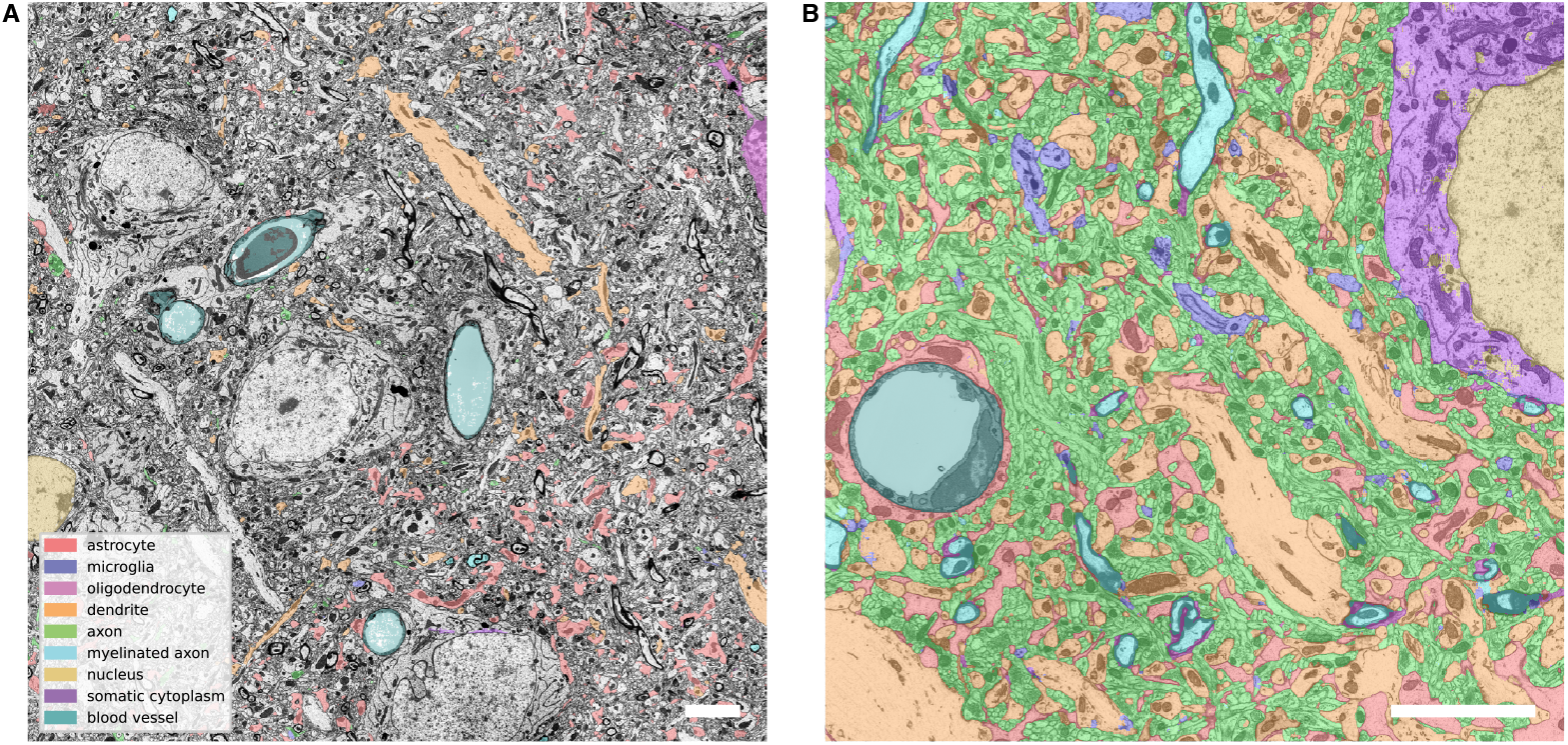
**A**: Ground truth annotation for semantic segmentation through instance segmentation supervoxel labeling. **B**: Sample model predictions illustrating all nine classes recognized by the model. Scale bar 5 µm.

## Materials and Methods

### Volume assembly

The individual tiles (sFOVs) acquired by the multibeam electron microscope were assembled using affine transforms into contiguous images of entire thin sections. These were then grouped into blocks of consecutive sections, with each block corresponding to a single 250 nm thick section. Block sizes ranged from 34 to 44 sections due to varying effective milling rates and the variability of physical section thickness.

We used SOFIMA (Scalable Optical Flow-based Image Montaging and Alignment) (Januszewski et al., 2024) to first align sections within each block, and then computationally flattened each block to a standard depth of 31 sections (corresponding to the 250 nm depth of thick sections before milling) with cubic interpolation in the, dimension. We used an intensity threshold of 70 to define a substrate mask for every section, which we then also dilated by 4 pixels. For every xy. position within each block we defined the valid tissue depth as the index of the first section within the block at which the substrate mask was active for the respective in-plane location. Only valid tissue data was interpolated, and pixels detected as substrate were excluded from the flattening process. We used images with intensity histograms matched to a reference distribution computed for a representative section, which was necessary to avoid „ringing” effects after flattening due to intensity distribution shifts between sections.

The thick sections were arbitrarily oriented when imaged by the microscope. We computed a global affine transform for every flattened block which brought them into approximate alignment. We then applied SOFIMA again over all blocks to build the final aligned volume (Supplementary).

### Semantic segmentation

We trained a neural network model to classify voxels into nine classes corresponding to selected cell types and compartments (Table S1). The model used image data at 32×32×32 nm^3^ voxel size and had a fully convolutional residual convstack architecture (He et al., 2016; Januszewski et al., 2018) (16 residual modules with 2 convolutional layers of 32 feature maps and 3×3×3 kernels each) with an effective field of view of 65×65×65 vx^3^ ≈ 2×2×2 µm^3^.

**Table S1.** Characteristics of the semantic segmentation training data.

| CLASS | SUPERVOXELS | POINTS |
| --- | --- | --- |
| astrocyte | 381 | 1,636,243 |
| microglia | 3 | 20,713 |
| oligodendrocyte | 232 | 5,532 |
| dendrite | 53 | 227,746 |
| unmyelinated axon | 282 | 81,521 |
| myelinated axon | 194 | 19,896 |
| nucleus | 152 | 32,218 |
| somatic cytoplasm | 203 | 21,928 |
| blood vessel | 8 | 126,046 |

We identified supervoxels representing different classes in an older instance segmentation of the volume (Table S1, Figure S1A). We then skeletonized that segmentation using the TEASAR algorithm (Sato et al., 2000), postprocessed to achieve an average node separation of 300 nm. During training, we sampled the centers of the training examples among the nodes of the skeletons corresponding to the annotated supervoxels, with all nine classes sampled with equal frequencies. The resulting model could reliably detect tissue types throughout the volume (Figure S1B).

### FFN (Flood-Filling Network) training

We enhanced the original FFN v1.0 architecture (Januszewski et al., 2018) and training scheme to increase segmentation accuracy, resulting in FFN v1.5. The overall convstack architecture was retained, but we extended the number of residual modules to 20, and introduced layer normalization (Lei Ba et al., 2016) after every residual module, which was necessary for the deeper models to train effectively. We also changed the optimizer to AdamW (Loshchilov et al., 2017) and used synchronous gradient accumulation during training. Training example centers were sampled uniformly at random among seed points in dendrites, axons and cell bodies within a densely segmented 8.8 × 8.8 × 26.4 µm^3^ subvolume (499 Mvx) at a voxel size of 16 × 16 × 16 nm^3^. The seed points were generated with the same procedure as used during inference (Januszewski et al., 2018). A subvolume shape elongated in the Z direction was chosen to increase the model’s exposure to thick section interfaces during training.

We used a densely skeletonized 12.3 × 12.3 × 12.3 µm^3^ subvolume to evaluate segmentations obtained with FFN checkpoints (snapshots of network weights) saved during training. We selected checkpoints corresponding to the highest edge accuracy (Januszewski et al., 2018) for full volume inference.

### FFN segmentation

We followed a coarse-to-fine segmentation strategy (Figure 1A) designed to minimize supervoxel-level merge errors while focusing computational cost to parts of the volume where it provides most added value. This was achieved by sequentially segmenting subsets (blood vessels, glia, everything else) of the volume identified by the semantic segmentation model and adjusting the number of inputs to oversegmentation consensus depending on the type of tissue being processed.

We started from a blood vessel segmentation generated at a reduced resolution of 32×32×32 nm^3^ per voxel, and segmented the volume independently with the top three FFN checkpoints, with the FFN FOV positions restricted to voxels labeled as astrocytes and microglia by the semantic segmentation model. We then combined these three segmentations with OC (oversegmentation consensus) (Januszewski et al., 2018), and removed any resulting supervoxels smaller than 10,000 voxels or for which microglia or astrocyte were not the dominant predicted voxel types.

Using the glia and blood vessel segmentation as the initial state, we performed six inference runs of the FFN with the top three FFN checkpoints (by edge accuracy), each time with seeds ordered by ascending and descending, coordinates. We then computed the OC of all these segmentations, with the supervoxels with a dominant classification as soma, nucleus or myelinated axons held back at the state corresponding to the segmentation with the top scoring checkpoint, with and without seed order consensus, respectively. The held back supervoxels were excluded from the rest of the consensus procedure because we observed they were unlikely to be involved in merge errors, due to both their spatially sparse distribution within the volume and large caliber relative to dendrites and unmyelinated axons. The reduction in merge errors achieved by OC outweighed the concomitant increase in splits by a factor of 5× (Figure 4A, Table 1).

We then skeletonized the resulting segmentation with TEASAR and computed a Voronoi partition of all segments using the skeleton nodes. We associated every Voronoi cell with the dominant voxel class (as predicted by the semantic segmentation model), built interface graphs for all cells of every base segment, and partitioned the graphs to separate the components with dominant myelin and nucleus/soma classification. The partitioned graph was treated as an agglomeration graph for the Voronoi cells and used to materialize a postprocessed base segmentation with myelinated axon fragments separated from the unmyelinated parts, and with neurites split off from their parent somas.

Finally, we applied FFN agglomeration (Januszewski et al., 2018) to all neurites with a dominant axon classification. We computed the agglomeration scores independently with the top three FFN checkpoints used for the base segmentation. The agglomeration graph was created from edges corresponding to segment pairs for which the scores generated by all three checkpoints fulfilled the standard agglomeration criteria (((*d*_*A*_ *<* 0.02 *d*_*B*_ *<* 0.02) (iou *>* 0.8 ^ *S*_*\*\**_ *>* 0.8) iou *>* 0.9 ^ *S*_*\*\**_ *S >* 0.9, see (Januszewski et al., 2018) for details).

We applied this segmentation workflow with FFN v1.0 and v1.5 models optimized with the same ground truth sets, which allowed us to quantify the impact of the architecture and training procedure improvements described in the previous section. We trained both models at an input resolution corresponding to the ground truth segmentation (16 × 16 × 16 nm^3^ per voxel), and used only a single checkpoint for FFN v1.0 agglomeration, where the ensembling of multiple checkpoints did not demonstrate a favorable split-merge error trade-off. We observed that relative to FFN v1.0, FFN v1.5 models reduce merge errors by 10 both in the base and agglomerated segmentation, while reducing splits respectively by 36% and 32%, and that a segmentation generated by a single forward pass of a FFN v1.5 model is already superior to all stages of FFN v1.0 segmentation (Table 1).

### Medium-context models: SENSE

SENSE models take as input a volumetric image and a „prompt” consisting of a small, spherically-masked fragment of a neuron segment located at the center of the input subvolume. This guides the model to segment the entire selected neuron within its FOV (Figure 2B).

To achieve high recall and complement the functionality of the FFN, we used a residual variant of the U-net architecture (Ronneberger et al., 2015) for the SENSE models, enabling a larger FOV than that processed by FFN’s convstack. Three variants of the network were trained: two with a 128×128×128 vx^3^ FOV, processing data at resolutions of 16×16×16 nm^3^ and 32×32×32 nm^3^ per voxel, and one with a 256×256×256 vx^3^ FOV processing data at 16×16×16 nm^3^.

The U-net architecture consists of a downsampling stack, a middle stack, an upsampling stack, and a final 3 × 3 × 3 convolution. The input is initially mapped to 8 feature maps using a pointwise convolution. Residual blocks consist of a sequence of: swish activation, convolution, swish activation, convolution, and an optional pointwise convolution to adjust the number of feature maps. Each residual block incorporates an additive skip connection to its input. The middle stack used two such residual blocks. The downsampling stack consisted of 5 (6 for the 256^3^ FOV model) superblocks, composed of three residual blocks followed by a swish activation, 2 × 2×2 average pooling with 2× 2 ×2 strides, a swish activation and another residual block. Each superblock also increased the number of feature maps at its input relative to the initial embedding by a factor of 1, 2, 4, 8, 8, respectively. Upsampling blocks started by concatenating their inputs to the output of the downsampling stack at matching spatial resolution. This data was then processed by a residual block, upsampled 2 × 2 × 2 with nearest neighbor interpolation and 2× feature map expansion, and processed by another residual block. All convolutional layers within the network utilize 3 × 3 × 3 kernels and the swish activation function unless stated otherwise.

The training data comprised 1,684 manually agglomerated axons randomly selected from the dataset. Binary segmentation masks of each axon were converted into probability maps for training: 0.95 for segment interior, 0.05 for segment exterior, and 0.5 for voxels outside the central spherical query region (radius of 16 voxels). The model was optimized using voxelwise cross-entropy loss to extend the prompt mask to the entire FOV in a single forward pass.

The probability map predicted by the SENSE model can be thresholded (here, at 0.5), and the results compared to the base segmentation. The *overlap score* for every base segment within the FOV of the model is then defined as the number of predicted positive (above threshold) voxels within that segment, divided by the number of voxels of that segment (original segment size).

We used a set of 320 axons (“validation set”) with a total path length of 54 mm for checkpoint selection. We identified all base segments corresponding to these axons and ran SENSE model inference with the FOV centered at every skeleton node within 2 µm of an endpoint (i.e., a node of degree one). For every inference call, we considered a positive prediction if the overlap score was at least 0.5, original segment size at least 500. If there were multiple positive predictions for a single inference call, we only retained the one with the highest overlap score.

We selected checkpoints in an iterative procedure, in which for every checkpoint we identified the true positive base segment pairs corresponding to the axons from the validation set. We then computed the number of connected components that would be formed, treating these pairs as edges in an agglomeration graph. The checkpoint corresponding to the lowest number of connected components was selected as the main checkpoint. Additional checkpoints were then added iteratively, at every step selecting a checkpoint that maximally reduced the number of connected components, given the edges associated with it and every previously selected checkpoint. This resulted in a total of seven checkpoints across the three model variants we trained achieving 97% recall over the validation axon set.

We then ran inference with these checkpoints over all axons within the volume, and identified agglomeration graph edge candidates as outlined above, allowing multiple edge candidates per SENSE inference call.

### Large-context models: SHAPE

To further filter the agglomeration candidates proposed by the SENSE models, we used SHAPE models with a spherical FOV 20 µm in diameter. The corresponding subvolume contains almost two gigavoxels of data and is too large to efficiently process with models that operate directly in voxel space. We therefore chose a point cloud representation, with the points sampled at random from faces of triangular meshes computed for the segments using marching cubes. SHAPE is a variant of the PointNeXt model (Qian et al., 2022) trained to detect whether point clouds represent plausible neurite shapes, i.e., are free of morphologically apparent merge errors.

We extracted points restricted to a sphere of radius 10 µm, sampling at an average density of 100 points per µm^2^ of neuron mesh surface. When restricted to the spherical region of interest, any mesh fragments disconnected from the central part were discarded and not sampled from. For training and inference, point sets were resampled to a standard size of 8192 points using farthest point sampling (for downsampling) or random point repetition (for upsampling)

Training examples were defined by a set of segments and the center location of the FOV sphere. We used pairs of segments and associated locations generated by incorrect SENSE model predictions as negative examples. Positive examples were sampled from correct predictions of the SENSE model, as well as directly from the skeleton nodes of the training axons. The training set was a superset of that used for the SENSE model and comprised 1,795 axon fragments, whereas the validation set comprised 263 axons. We selected the model with the highest *F*_1_ score computed over the validation set for inference.

We explored various hyperparameters of the model and found that the highest *F*_1_ score was achieved by a variant trained with *n* = 16 independent point cloud samples per example, 8192 input points, and using *k* = 16 nearest neighbors in each of the six downsampling stages within the network.

We used the checkpoint corresponding to the maximum *F*_1_ score to evaluate all pairs of segments generated by the SENSE models. We considered segment pairs with overlap scores for both segments of at least 0.5, original segment size at least 500 voxels at the model input resolution, and for which at least one segment had a skeleton endpoint within the FOV of the model. Blood vessels, segments with the top three semantic model classifications other than axon and larger than 20,000 voxels, as well as segments classified as astrocytes or dendrites were excluded from the SENSE results. We also excluded segments with less than 2500 nm path length. For every remaining segment pair, we used all skeleton nodes of either segment that were within 2 µm of the SENSE FOV center for SHAPE inference.

### SHAPE: Scaling

We observed that averaging SHAPE predictions over random rotations and point cloud samples each improve accuracy (Figure S2B,C). Their combined use produces a synergistic effect, yielding gains that exceed their individual contributions (Figure S2C). This stands in contrast to earlier stages of PATHFINDER, where ensembling FFN and SENSE models allows for trade-offs between recall and precision, but not simultaneous improvements in both (Figure 4). SHAPE is the only stage of the pipeline where additional inference-time compute, without changing how the model is applied, translates directly into higher quality results, reducing the *F*_1_ error by up to 44% (Figure S2B). For the results presented in this work, we averaged over 16 point cloud samples, each with 16 random rotations.

The additional compute cost that this incurs might in the future be partially mitigated by using rotationally invariant representations (Unke and Maennel, 2024), and different point sampling schemes. Out of the two, point sampling is the dimension along which the SHAPE models currently exhibit higher variance (Figure S2A).

**Figure S2.**
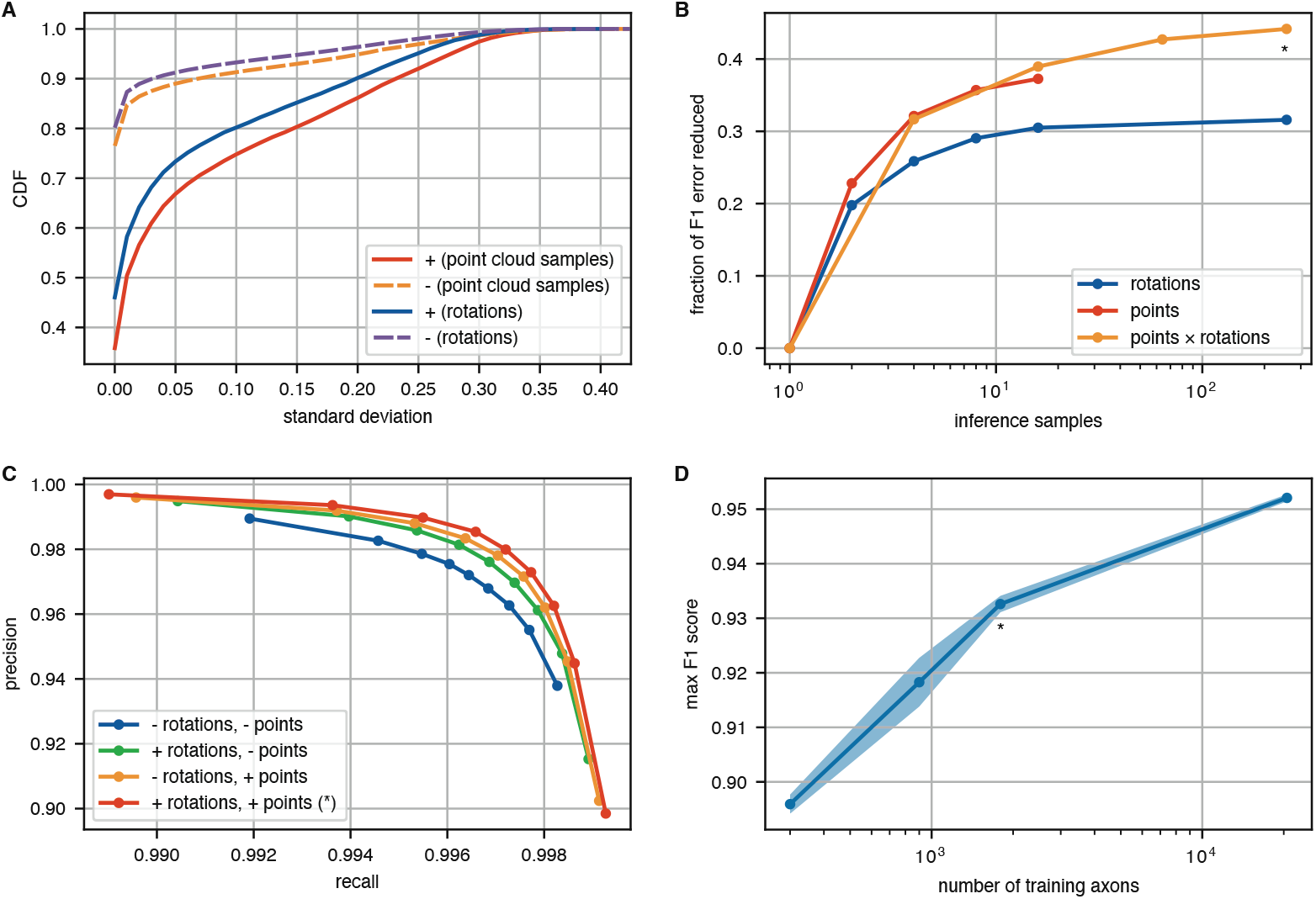
Impact of inference time (A-C) and training set (D) scaling on the performance of false positive (false merger, -) axon segment pair detection with SHAPE (*r* = 10 µm, 8192 points). **A**: Cumulative distribution (over the test set) of standard deviation of classifier predictions due to rotations and point cloud sampling. **B**: Reduction in *F*_1_ error (1 − *F*_1_) of the base model by averaging over random rotations and point cloud samples for every evaluated segment pair. **C**: Precision-recall curves with different combinations of rotation and point cloud sampling averaging. **D**: *F*_1_ score as a function of training set size for SHAPE models. Shaded areas indicate ± 2 standard errors (over top ten network checkpoints). B-C show evaluations for all positive and negative examples involving axons from the test set. D shows evaluation computed for an evenly weighted subset of positive and negative examples. * marks the settings used in the experiments reported in the paper.

We also observed that SHAPE accuracy improves logarithmically with increasing training dataset size (Figure S2D). We experimented with training on ∼ 50% unproofread axons in the volume and as shown in Figure S2D, the logarithmic scaling law might not be maintained in that regime. Larger volumes are needed to conclusively evaluate the limitations.

### SHAPE: Locations affiected by dust particles

We noticed a concentration of split errors (Figure S3C) in the direct vicinity of dust particles, which can obscure the view throughout most of the depth of the thick section (Figure S3B). These locations were also associated with a lower precision of the SENSE models, which together caused many proposed agglomeration graph edges to be generated in a spatially compact area. Observing that most of the splits separate axons into two pieces, one “above” the dust particle, and one “below”, we elected to preprocess the candidate agglomeration graph to take advantage of this insight.

**Figure S3.**
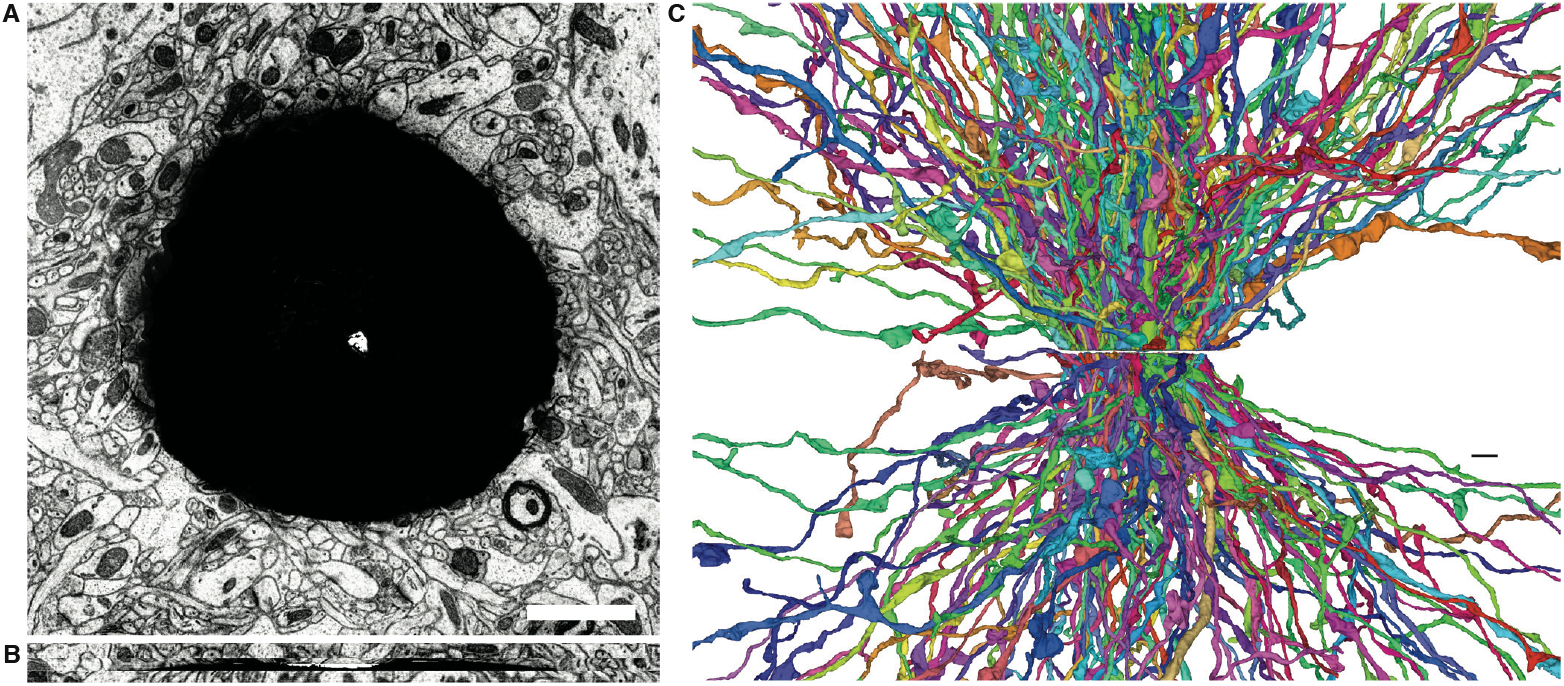
Dust particles and their impact on EM images and segmentation. **A**: XY view of a dust particle artifact. **B**: XZ view of the same particle. A single thick section is affected throughout its complete depth (250 nm), while adjacent sections are unperturbed. **C**: Dust particle causing splits in the axons that intersect the affected image area. Both scale bars 2 µm.

We processed every dust particle separately by extracting a subvolume fully containing it according to Table S2. Within the subvolume, we filtered the EM image of the last section (where the particle was most extensively visible) with a uniform filter of kernel size 10 and took voxels with intensity lower than 20 or higher than 240 in the filtered image as an artifact mask. We then computed the connected components of this mask, removed components smaller than 1,000 voxels, and dilated the mask with a radius of 10. Any segment intersecting that mask within 15 sections directly above or below the particle subvolume was considered affected by the artifact. For these segments, we computed the local connected components, and identified the range of, coordinates they span. Any component present 10 sections above the particle was considered a member of the “above” set, and any component present 10 sections below it was considered a member of the “below” set. Segments present in both sets were considered as successfully crossing the artifact.

We identified 357 axons affected by seven of the eight dust particle artifacts present in the volume (Table S2). One particle was excluded from processing due to its location being directly adjacent to the volume boundary, which did not provide sufficient context for SENSE and SHAPE models. SENSE model recall for the affected axons was 95.2%.

For every particle, we modeled the local agglomeration decisions as a bipartite matching problem between the “above” and “below” segments. Specifically, we used the SHAPE scores as weights in a linear sum assignment problem, which we solved with the Jonker-Volgenant algorithm. Applying a SHAPE score threshold of 0.9 for filtering resulted in 238 true positive and 19 false positive agglomeration decisions, yielding a recall of 67% and a precision of 93%.

### SHAPE: Combinatorial search

To take advantage of SHAPE’s ability to evaluate arbitrary reconstructions irrespective of their supervoxel composition, we applied the model to objects corresponding to higher order motifs of the agglomeration graph. We started with an initial graph formed from SENSE predictions and filtered by pairwise SHAPE predictions with a mean score of at least 0.5 (Figure 4A). This corresponded to a setting with few split errors and many merge errors, the latter of which we intended to reduce while incurring a minimal number of additional splits.

We considered a 1 × 1× 1 µm^3^ grid of points covering the volume, and defined two subvolumes centered at every point: a 1 × 1 × 1 µm^3^ „core” subvolume and a 10 10 10 µm^3^ „context” subvolume. For every point, we computed the connected components of the prefiltered agglomeration graph restricted to the context subvolume, and selected for further processing only those components that had at least one edge located within the core subvolume. We then used SHAPE to evaluate objects formed by connected subgraphs of these components up to order four.

Any subgraph scored lower than a threshold value was considered invalid. For every grid point, and for every component we then solved an optimization problem aiming at identifying a set of agglomeration graph edges that need to be removed so that the resulting graph minimizes the number of connected components (i.e., few new split errors are introduced), while eliminating all invalid subgraphs (i.e., any identified merge error is eliminated). We did this in an iterative procedure, in which the nodes of every component start completely separated. The original component edges are then processed in order of descending pairwise SENSE (more mergers, fewer splits) or SHAPE (fewer mergers, more splits) scores and added to the graph as long as no invalid subgraph is formed. The edges removed from the original agglomeration graph is then considered the „solution” for that specific point and component.

Following the generation of initial per-point, per-component solutions, a global reconciliation procedure was performed. First, solutions derived for specific invalid subgraphs were evaluated for applicability to all other points and components associated with the same subgraph. The global occurrence frequency of each distinct solution was then tabulated. Finally, each point and component was revisited and assigned the solution exhibiting the highest global frequency among its applicable candidates.

Figure 4A shows the Pareto frontier of results obtained with this procedure, which is significantly better than pairwise filtering with SHAPE alone. The total number of errors was minimized by setting the SHAPE decision threshold at 0.2.

### SHAPE: Merge error detection

SHAPE can identify and localize parts of segments with suspected merge errors. This could be used to target human attention to specific cases that need inspection and potential corrections. We checked if this presents a viable approach for identifying merged supervoxels in the base segmentation. This is a challenging problem since these errors are quite rare (Figure 4A).

Application of SHAPE to all axons within the base segmentation identified 307 segments containing connected skeleton components of at least 20 nodes (approx. 6 µm linear extent), for which SHAPE flagged every constituent node. SHAPE predictions were considered to indicate a merge error at the associated location when the standard deviation over 16 random rotations was lower than 0.1, and when the mean prediction over rotations and point cloud samples was lower than 0.1.

A review of these results showed 30 actual merge errors, most involving small fragments merged into the axons. 7 of these had a sufficiently large spatial footprint to be reported in the standard skeleton-based evaluations. These global metrics also identified 28 other merge errors that were not detected as such by SHAPE. Of these, 5 were flagged by SHAPE but involved less than 20 points or the flagged points did not form a connected component. The remaining 23 involved „jumps” to an incorrect axon while tracing through a thick section interface. These mistraced axons appear locally morphologically correct, and the associated errors can only be detected when looking at the premature terminations (split errors) of the correct continuations of both axons involved in the merger.

Observation revealed that in addition to identifying genuine merge errors, SHAPE also flagged locations exhibiting unusual yet correctly reconstructed morphologies. Examples of such instances included axons with numerous small branches, self-contacting branches, and branches invaginating adjacent neurites.

## Supplementary Text

### Segmentation evaluation

We used topological, skeleton-based metrics (Januszewski et al., 2018) to automatically evaluate the quality of the different stages of reconstruction (Table 1, Figure 4). We skeletonized a reference volumetric segmentation of the complete volume with the TEASAR algorithm (invalidation scale 5, invalidation constant 0). We then postprocessed the skeletons by eroding endpoints by 100 nm and maintaining an average node separation of 300 nm.

We used all axons in the thus created ground truth skeleton set to compute splits and merge count and nERL (normalized expected run length). The number of split errors associated with each reference skeleton was the number of segments that overlapped it over at least five nodes, reduced by one. For this count we did not include segments that did not extend beyond 144 pixels away from a face of the volume bounding box, for which insufficient spatial context was present for the large FOV models to effectively agglomerate them. The number of merge errors associated with each reconstructed segment was the number of distinct skeletons overlapping that segment over at least 10 nodes, reduced by one. The node count thresholds were used to make the metrics insensitive to minor errors in the ground truth set, as well as minor merge and split errors that do not affect the global topology of the neurons.

To focus analysis on the most challenging tracing problems, we restricted evaluation to unmyelinated axons, of which the total path length in the volume was 4,284 mm. Myelinated axons have calibers comparable to dendrites and are therefore significantly easier to trace, a task further simplified by the presence of the myelin sheath with its distinct appearance in EM images.

### Volume statistics

The total analyzed volume is a cuboid 80.4 × 84.8 × 99.2 µm^3^ = 676,338 µm^3^ in size, itself inscribed into a slightly larger and irregular block of tissue.

We proofread the complete volume by screening all axon fragments longer than 5 µm for false mergers and splits based on morphological cues, and selectively followed through with an examination of the corresponding EM image content to resolve any identified putative errors.

After proofreading, over 59,000 axon fragments with a total path length of 4.45 m were identified within the volume. Not all of these fragments have pre-synaptic sites, so a small fraction might belong to another category (microglia, oligodendrocyte, aspiny dendrite). Similarly, some short axon fragments might have rarely been mistakenly identified as belonging to another class.

### Dust particles

We identified eight instances of dust particles embedded between the section and the wafer (see Figure S3A for impact on EM images and Table S2 for characteristics of location and extent). These particles completely obscure the tissue content within an area of up to a few square microns and over a depth of up to the full thickness of the section before milling (here,∼250 nm). Despite this missing image information, we found that a human expert can in all cases connect neuron fragments across these particles, even if this process requires significantly more mental effort than tracing neurons in unaffected areas.

The genesis of these artifacts is well understood, and they can be eliminated or at least significantly reduced by strict environmental control during wafer preparation.

**Table S2.** Approximate location and extent of the artifacts caused by the presence of dust particles between the section and the wafer.

| x [vx] | y [vx] | z [vx] | r [μm] | d [sections] |
| --- | --- | --- | --- | --- |
| 5878 | 5050 | 4416 | 2.5 | 12 |
| 4274 | 1962 | 433 | 2.5 | 16 |
| 2571 | 3836 | 1115 | 2.5 | 16 |
| 3318 | 4411 | 2324 | 5.0 | 16 |
| 5902 | 5443 | 2975 | 2.5 | 6 |
| 5412 | 2410 | 3161 | 2.5 | 14 |
| 5777 | 1818 | 4540 | 4.0 | 15 |
| 990 | 3913 | 5982 | 3.0 | 16 |

### Analysis of reconstruction errors

We skeletonized the segments involved in the 2,816 unique splits (some splits out of the 2,862 total are double counted because of merge errors) and identified their endpoints (nodes of degree one). For every segment we then counted the number of endpoints located within a „margin area” defined as locations up to 64 voxels away from a face of the bounding box of the volume. This allowed us to classify segments as orphans (no endpoints in the margin areas), single-face-crossing (SFC, endpoints within the margin area of a single face) and multi-face-crossing (MFC). Among the axon segments, we found 9,324 SFC with one endpoint in the margin area, 6,883 SFC with more than one endpoint in the margin area, and 2,265 orphans. 1,668 and 361 of the SFC segments with one or more endpoints, respectively, were involved in a split error.

We found 59 instances (2% of all false splits) where the split itself was located in the margin area, 1,354 (48%) instances involving orphan segments, and 1,638 (58%) instances involving SFC segments. Overall, 2,517 (89%) false splits involved either an orphan or an SFC segment.

542 false splits (19%) involved a segment affected by a false merge error, 403 of which also involved either an orphan or an SFC segment. A further 219 false splits involved segments with minor mergers (less than 10 skeleton nodes). These mergers each cause at most one synapse to be falsely assigned and are thus comparable to a dendrite spine-trunk misassignment in terms of their impact on the connectome matrix. If we consider this acceptable and retain them in the final reconstruction, 194 of the related false splits do not need to be addressed, leaving only 25 splits that do (all of which involve orphan or SFC segments). Overall, just 141 (5%) of splits involved only MFC segments, and in all these cases at least one of the segments was affected by a merge error.

### Proofreading cost estimates

For comparison with prior estimates, we use „reconstructed volume per time” or PT (proofreading throughput) as a universal measure of reconstruction efficiency. Real-life proofreading workflows are often organized into different phases (e.g. elimination of large-scale mergers, connecting large segments to a soma, etc (Nern et al., 2025)). PT abstracts away from these details and represents an average over all activities that are necessary for a volume of tissue to be considered completely reconstructed.

The human labor component of a whole mouse brain connectome project has recently been estimated at $6B-$20B, corresponding to 0.3-1 million person-years of work (Jefferis et al., 2023). The volume of a mouse brain was assumed to be ∼500 mm^3^ and therefore with a person-year of 2080 h, the shorter end of the estimate corresponds to a PT of 800 µm^3^/h.

To estimate an upper bound of PT for the present study, we consider the time necessary to fix all errors affecting axons within the volume. As explained in the main text and noted in prior work (MI-CrONS Consortium), with their higher caliber trunks and lower total path length, dendrites represent a small fraction of effort needed for axons. This results in a PATHFINDER PT of 67 200 µm^3^/ h, assuming 10 s as a optimistic mean time to fix a split or merge error. In our experience, this amount of time is sufficient to manually identify and select a partner segment that will correctly continue to trace a starting segment provided the cursor of the viewer is pre-positioned in the vicinity of the split location. This time could be further minimized if candidate segments are already available (e.g. preselected by SENSE) and the tracing could be reduced to one or more binary (accept/reject) decisions. In the case of mergers, human proofreaders exhibit excellent intuition about which shapes should be separated, and modern user interfaces make it possible to do so by placing seeds in the objects that need to be split (Dorkenwald et al., 2025; Hubbard et al., 2020).

In order to be efficiently solved, errors also need to be detected and localized. In sufficiently large volumes, false mergers eventually lead to violation of known biological priors, such as multiple somas connected to a single segment. SHAPE models can also be used to flag putative merge errors for human review as discussed previously. Similarly, split errors in large volumes cause segments to remain „orphans” (fully contained within the volume and not connected to a soma).

In smaller volumes, prioritizing the review of orphans and SFC segments is an effective strategy for identifying false splits. In the present volume, as outlined in the previous section, this approach makes it possible to detect 95% of all false splits. Only a subset of all SFC segments are involved in split errors, but the presence of a synapse at the endpoint furthest away from the face of the bounding box of the volume that the segment crosses can be an effective filter. To show this, we manually inspected 100 randomly selected SFC segments, and found 21 to be involved in a split error. Of these, 16 did not terminate at a synapse. 5 did, but connected to another SFC or orphan segment. Of the remaining 79 segments, all terminated at a synapse.

In prior estimates (Jefferis et al., 2023), the proofreading effort for a whole mouse connectome was projected to last 10 years. At the originally estimated speeds, this would require an unrealistically large workforce of at least 30,000 full-time proofreaders. PATHFINDER reduces this by a factor of up to 84×, making it only an order of magnitude larger than previously completed brain mapping projects in Drosophila (Dorkenwald et al., 2024; Scheffer et al., 2020).

